# Reads2Resistome: An adaptable and high-throughput whole-genome sequencing pipeline for bacterial resistome characterization

**DOI:** 10.1101/2020.05.18.102715

**Authors:** Reed Woyda, Adelumola Oladeinde, Zaid Abdo

**Affiliations:** Department of Microbiology, Immunology and Pathology, Colorado State University, Fort Collins, CO, USA; Program in Cell and Molecular Biology, Colorado State University, Fort Collins, CO, USA; Bacterial Epidemiology and Antimicrobial Resistance Research, USDA-Agricultural Research Service, Athens, Georgia, USA

## Abstract

**Summary:** The bacterial resistome is the collection of all the antibiotic resistance genes, virulence genes, and other resistance elements within a bacterial isolate genome including plasmids and bacteriophage regions. Accurately characterizing the resistome is crucial for prevention and mitigation of emerging antibiotic resistance threats to animal and human health. Reads2Resistome is a tool which allows researchers to assemble and annotate bacterial genomes using long or short read sequencing technologies or both in a hybrid approach. Using a massively parallel analysis pipeline, Reads2Resistome performs assembly, annotation and resistome characterization with the goal of producing an accurate and comprehensive description of a bacterial genome and resistome contents. Key features of the Reads2Resistome pipeline include quality control of input sequencing reads, genome assembly, genome annotation, resistome characterization and alignment. All prerequisite dependencies come packaged together in a single suit which can easily be downloaded and run on Linux and Mac operating systems.

**Availability:** Reads2Resistome is freely available as an open-source package under the MIT license, and can be downloaded via GitHub (https://github.com/BioRRW/Reads2Resistome).

## 1. Introduction

Antimicrobial resistance is on the rise worldwide and the CDC estimates that per year at least two million people will become infected by a drug-resistant bacteria and at least 23,000 will die as a result of such infections [1]. The primary method for determining antimicrobial resistance in clinical laboratories is culture-based antimicrobial susceptibility testing (AST). However, the declining cost of next-generation sequencing technologies has enabled increased sequencing depth and more accurate identification of antimicrobial resistance elements within bacterial genomes [2]. Additionally, PacBio and Oxford Nanopore MinION sequencing technologies provide sequencing reads longer than 10kb, improving our ability to generate complete and accurate genome assemblies. This has led to an increase in open-source bioinformatics tools which can leverage multiple types of sequencing data. When performing an analysis such as identifying antimicrobial resistance genes, a researcher might use a dozen or more individual tools with each relying on their own databases and dependencies. Here we present Reads2Resistome, a streamlined bioinformatics pipeline for quality control, assembly and resistome characterization for bacterial genomes which analyzes short reads, long reads or both in a hybrid assembly approach.

## 2. Methods and Implementation

Reads2Resistome is scripted using Nextflow [3], a parallel DSL workflow framework, and is integrated with Singularity [4], an open source container platform with focus towards HPC workloads. The Reads2Resistome pipeline includes three main steps: quality control, assembly, and annotation of input bacterial sequencing reads. Reads2Resistome takes long read sequences, short read sequences or both as input and performs quality control of short reads using Trimmomatic [5] and long read quality visualization using NanoPlot [6]. Both quality-controlled short reads and long reads are then assembled using Unicycler [7], which generates an assembly graph using short reads, then uses long reads to simplify the graph to generate accurate assemblies. In the event the input consists of short reads only, Unicycler employs SPAdes [8,9] for assembly and subsequently polishes the resulting graph by bridging contigs. Long read-only assembly is performed using miniasm [10] and Racon [11] employed through Unicycler. Annotation is performed with Prokka [12] using one of the provided custom databases, which are pre-built from collections of specific bacterial species and subtypes, or using the Prokka default database. Resistome characterization is performed using ABRICATE [13]. Nextflow implementation using Singularity gives version control over the various open source tools ensuring reproducible results. Reads2Resistome output contains the following for each input isolate: visualization of both raw and quality-controlled reads; assembled contigs with a corresponding assembly graph along with an assembly quality assessment; gene and resistome annotation files; genome alignment files in BAM format; and optional serovar predictions. All documentation and pipeline usage is publicly available at https://github.com/BioRRW/Reads2Resistome.

### 2.1 Streamline high-throughput analysis

Reads2Resistome is designed for high-throughput bacterial sequence input and performs quality control, genome assembly and subsequent genome and resistome annotation in a parallel, high-throughput manner (**Figure 1**). Reads2Resistome is able to accommodate input of different species within the same run and can perform species-specific genome assembly quality control and gene annotation. A comma-separated values (CSV) file, generated by the user, enables input of multiple different isolates regardless of the isolate identity. The user can also specify a pipeline-provided database for genome annotation or can choose to use the default database utilized by Prokka. For genome assembly quality assessment, done by QUAST, the user can optionally add a user-provided reference genome for additional reference-specific metrics. Pipeline outputs for quality control and genome quality assessment are aggregated by MultiQC into a HTML report. In addition to quality-control, assembly and annotation, genome alignments are generated for further comparison. For *Salmonella spp*. optional serovar prediction is performed using SISTR.

**Figure 1.**
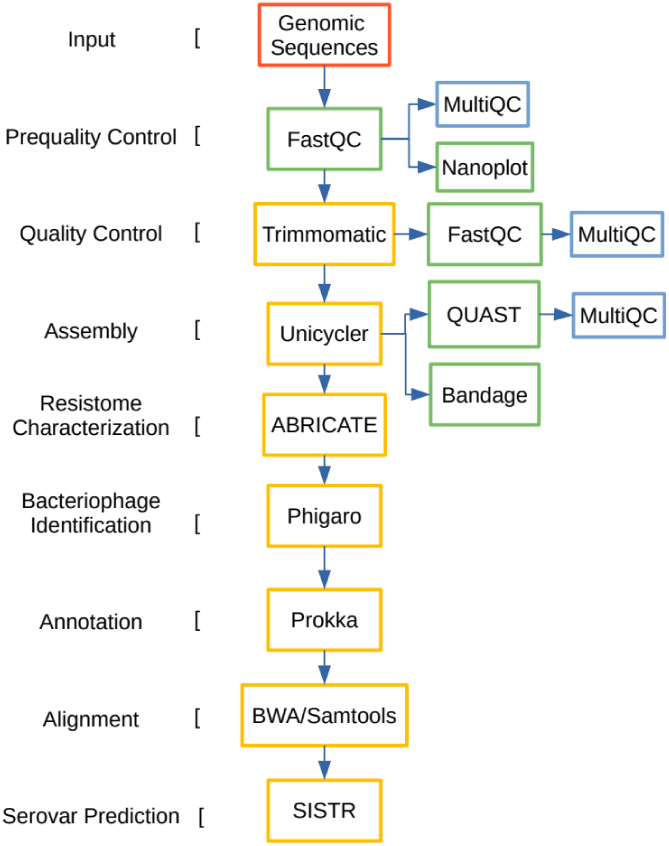
Key processes in Reads2Resistome pipeline.

### 2.2 Adaptable to cutting-edge sequencing technologies

Inclusion of long read sequences into bacteria assembly aids in resolving repeat regions of genomes and contributes to genome completeness [14]. Reads2Resistome is designed to be adaptable and flexible in that it can accommodate assembly in three different approaches: long read-only assembly, short read-only assembly, and hybrid assembly. In each assembly approach, Unicycler is used to generate genome assemblies and assembly graphs which are visualized with Bandage (**Figure 1**) [15].

### 2.3 Resistome characterization

Reads2Resistome characterizes resistome contents using ABRICATE and Phigaro. ABRICATE uses the assembled contigs to screen for antimicrobial and virulence genes from various databases; ARG-ANNOT antibiotic resistance gene database [16], the Comprehensive Antibiotic Resistance Database (CARD) [17], MEGARes Antimicrobial Database for High-Throughput Sequencing [18], NCBI AMRFinderPlus [19], PlasmidFinder [20], ResFinder [21] and VirulenceFinder database [22]. ABRICATE compiles results into a single report containing hits from each database and Reads2Resistome provides an output file for each isolate. Phigaro uses the assembled contigs to detect putative taxonomic annotations and the output is collected by Reads2Resistome for each isolate.

## 3. Case Study

Using Reads2Resistome we assembled and characterized the resistome of two bacterial isolates recovered from the ceca of 2-week old broiler chickens; SH-IC: *Salmonella enterica* serovar Heidelberg (S. Heidelberg) and EC-IC: *Escherichia coli*. (**Table 1**). Illumina, PacBio and Oxford Nanopore MinION sequences were used to test each type of assembly available under the pipeline; short read-only, long read-only and hybrid assembly.

**Table 1.** Summary of isolate information.

| Bacterial strains | Strain ID | SRA accession no. | Sequencing platform | Coverage* |
| --- | --- | --- | --- | --- |
| *Escherichia coli* | EC-IC | SRR11808523 | Illumina | 56.95 |
|  |  | SRR11808522 | MinION | 35.35 |
|  |  | SRR11808521 | PacBio | 28.08 |
| *Salmonella* Heidelberg | SH-IC | SRR11808520 | Illumina | 20.25 |
|  |  | SRR11808519 | MinION | 16.69 |
|  |  | SRR11808518 | PacBio | 33.14 |

\* Coverage estimated from total quality-controlled bases divided by the genome size (*E. coli*: 4800000bp, *S. heidelberg*: 4600000bp)

Short and long read-only assemblies, regardless of the read source, resulted in the shortest run-time with an average of 6 minutes per sample. Hybrid assembly, as expected, was the most time-intensive assembly method taking on average 1 hour and 8 minutes per sample, regardless of long read source (**Table 2**).

**Table 2.** Summary of assembly resources and run-time under various assembly conditions.

| Isolates included in run | Assembly Method | Elapsed time | --threads (option) | CPU-hours |
| --- | --- | --- | --- | --- |
| EC-IC Illumina;<br>SH-IC Illumina | Short Read | 16m 13s | 64 | 0.6 |
| EC-IC MinION;<br>EC-IC PacBio;<br>SH-IC MinION;<br>SH-IC PacBio | Hybrid | 4h 31m 45s | 64 | 5.3 |
| EC-IC MinION;<br>EC-IC PacBio;<br>SH-IC MinION;<br>SH-IC PacBio | Long Read | 23m 46s | 64 | 0.9 |

Genome assembly and annotation metrics were compiled from QUAST and Prokka outputs. Hybrid assembly of both EC-IC and SH-IC using MinION long reads and Illumina short reads gave the fewest contigs, longest total length and highest number of annotated genes as compared to long read assembly using MinION. Hybrid assembly of both isolates using PacBio reads resulted in fewer contigs but comparable total length to that of the MinION hybrid assembly. Genome contiguity was best obtained by hybrid assembly and can be visualized with Bandage-generated graphs (**Table 4**). While hybrid and long read-only assemblies are comparable with respect to number of contigs and genome length, the long read-only assembly greatly lacked in annotated genomic features and resistome elements.

Annotated genes and features across all assembly methods for both isolates were considerably reduced under the long read-only assembly, while both short read and hybrid methods resulted in comparable numbers of annotated genes. We suspect this is due to relative lower quality of long reads as compared to Illumina short reads. (**Table 3**). This is mirrored in resistome characterization and bacteriophage identification. While both short read and hybrid assembly methods for both isolates resulted in comparable identified resistome elements and bacteriophages, long read-only assembly identified elements were significantly reduced (**Table 5**,**6**).

**Table 3.**
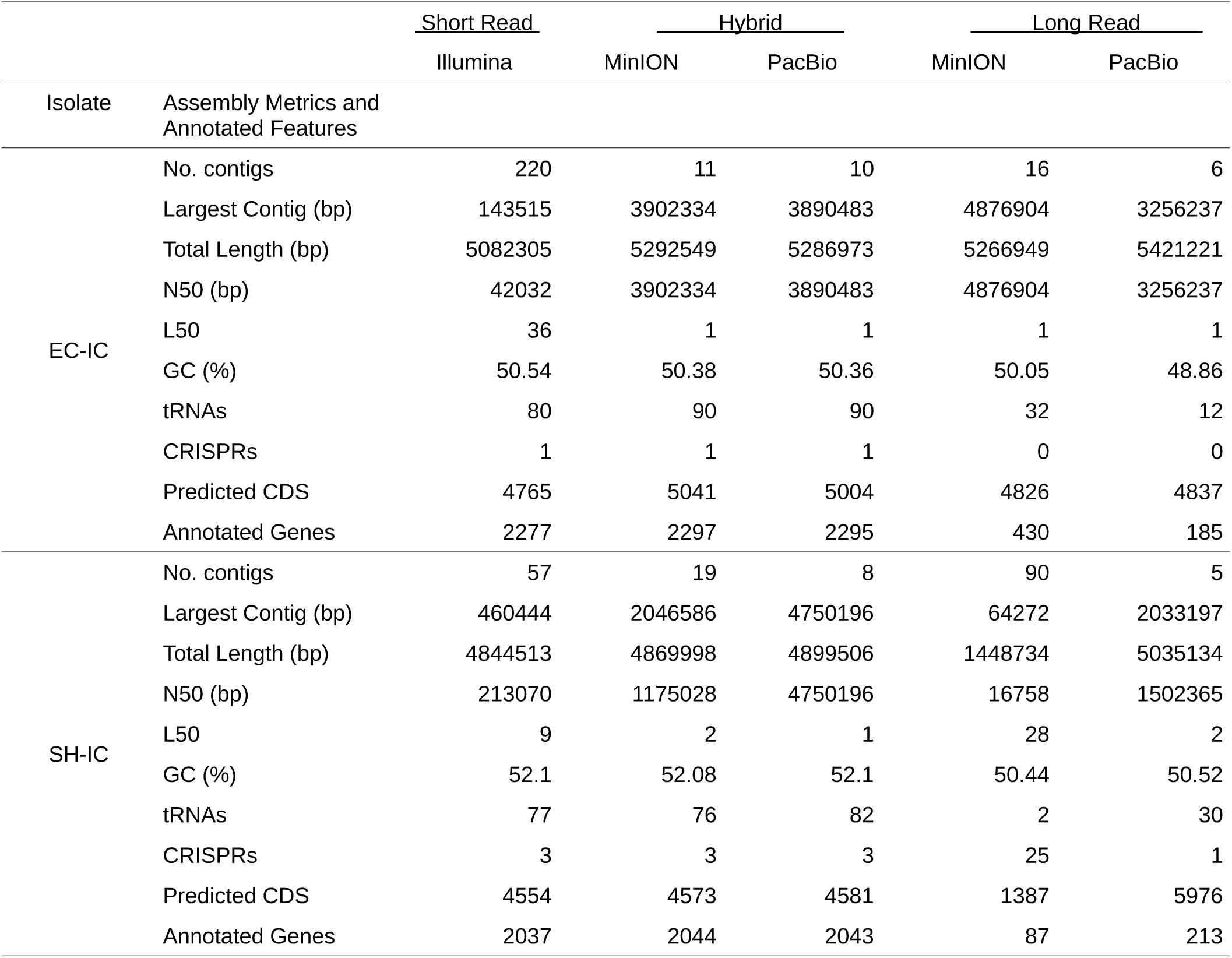
Summary of evaluation for assembled isolates under various assembly conditions.

**Table 4.**
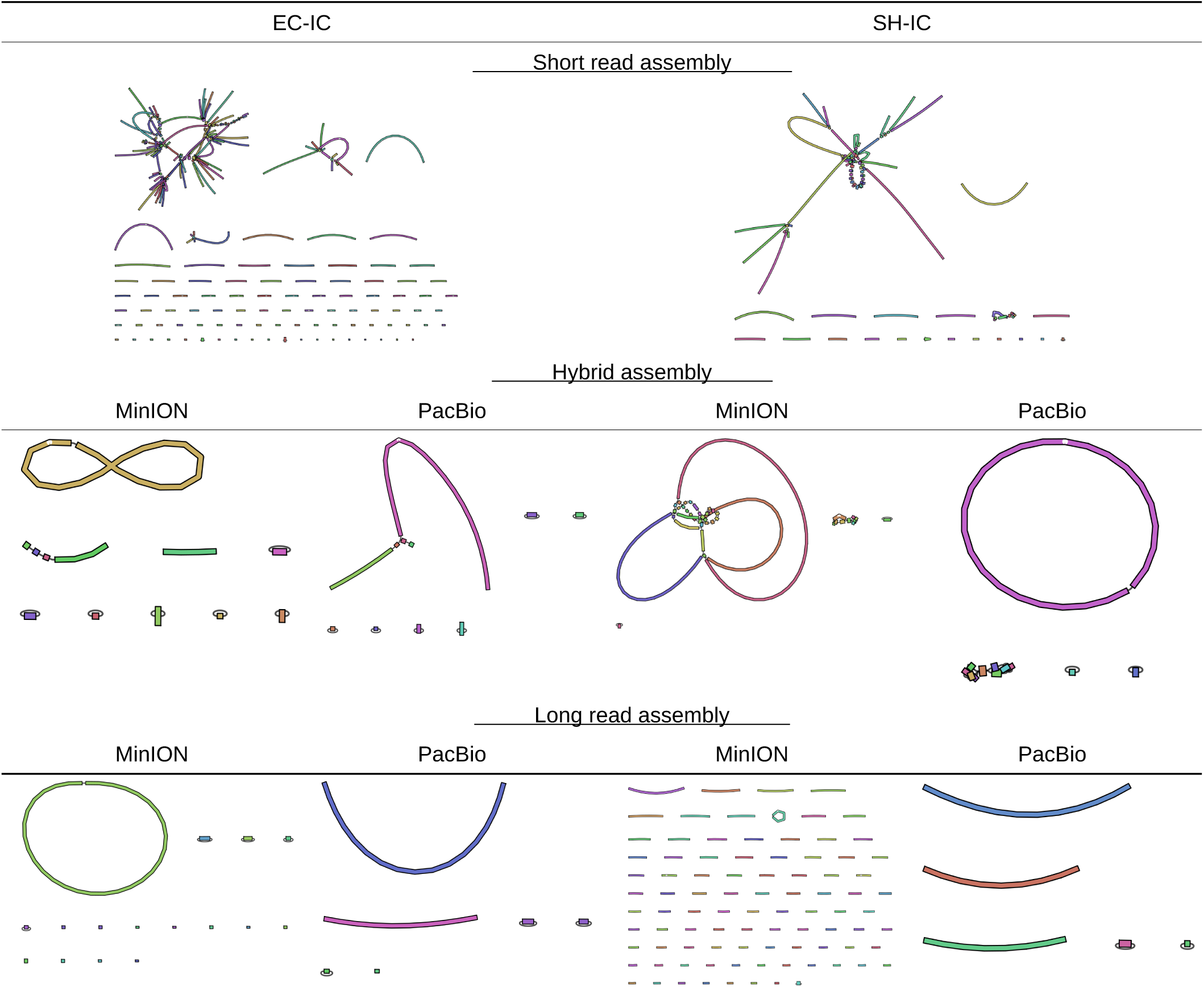
Bandage graphs for isolates under various assembly conditions.

**Table 5.** Summary of resistome characterization for assembled isolates under various assembly conditions.

|  |  | Short Read | Hybrid |  | Long Read |  |
| --- | --- | --- | --- | --- | --- | --- |
|  |  | Illumina | MinION | PacBio | MinION | PacBio |
| Isolate | Database | Number of identified elements |  |  |  |  |
| EC-IC | ARG-ANNOT | 8 | 8 | 8 | 7 | 8 |
|  | CARD | 48 | 48 | 48 | 44 | 43 |
|  | MEGARes | 58 | 58 | 58 | 53 | 52 |
|  | NCBI | 4 | 4 | 4 | 3 | 4 |
|  | PlasmidFinder | 7 | 7 | 7 | 5 | 5 |
|  | ResFinder | 4 | 4 | 4 | 3 | 4 |
|  | VirulenceFinder | 70 | 72 | 72 | 55 | 55 |
| SH-IC | ARG-ANNOT | 8 | 8 | 8 | 6 | 5 |
|  | CARD | 29 | 30 | 30 | 6 | 9 |
|  | MEGARes | 36 | 37 | 37 | 6 | 16 |
|  | NCBI | 4 | 4 | 4 | 4 | 3 |
|  | PlasmidFinder | 3 | 3 | 3 | 2 | 2 |
|  | ResFinder | 5 | 5 | 5 | 5 | 4 |
|  | VirulenceFinder | 105 | 105 | 106 | 44 | 99 |

**Table 6.**
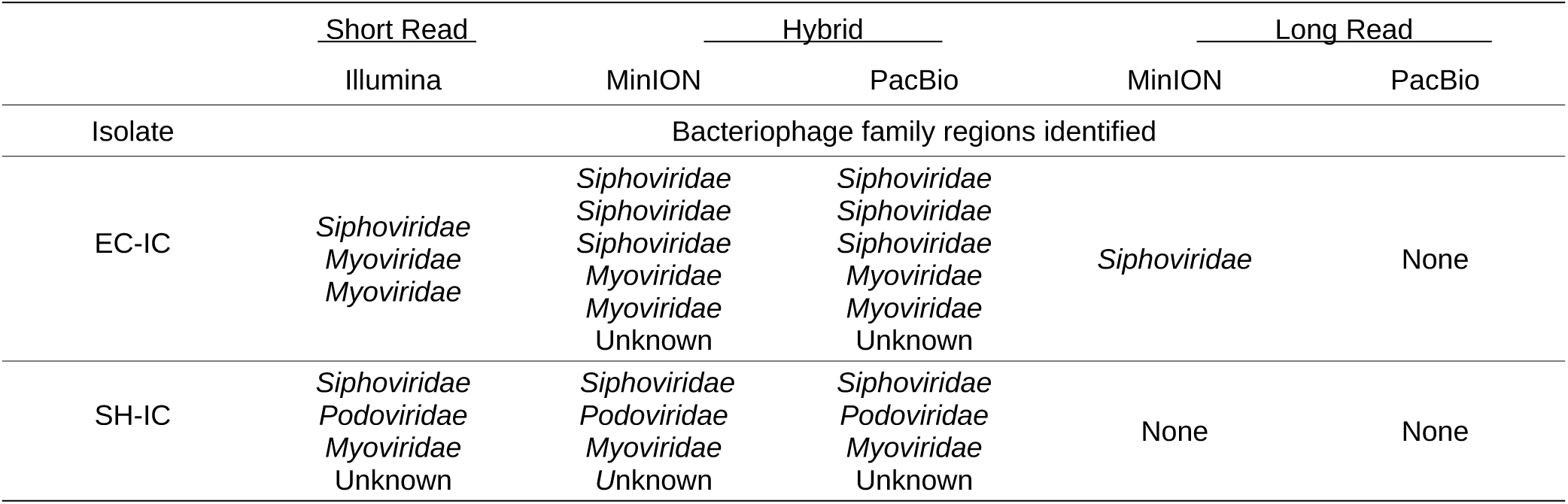
Summary of identified bacteriophages under various assembly conditions.

The pipeline was run using the following commands for short read-only, hybrid, and long read-only:

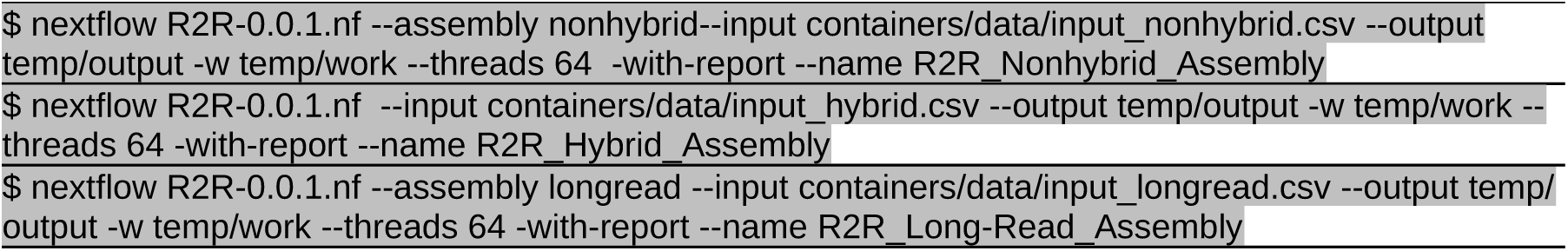

Each command was executed independently on a Linux server with 128 compute cores and 504GB of memory. Resources allocated and run-time in **Table 2** were obtained from the “report.html” which is generated using the ‘-with-report’ option.

## 4. Limitations

Reads2Resistome is designed to be deployed on Linux servers with at least 16GB of RAM and at least 16 compute cores. Running Reads2Resistome on a personal machine will allow full completion of the pipeline but time requirements will be daunting using less than 16 compute cores. Prokka gene annotation custom databases consist of *Escherichia coli, Campylobacter, Salmonella, Enterococcus and Staphylococcus*; all other input samples will use the default Prokka database. Currently serovar prediction is only provided for *Salmonella spp*. using SISTR. Additionally, obtaining high-quality genome assemblies along with accurate resistome characterization is dependent on the quality and depth of sequencing obtained for the input isolates.

## Conclusion

Reads2Resistome provides a streamline high-throughput analysis pipeline for the assembly, genome annotation and resistome characterization for bacterial sequenced reads. The pipeline can perform three methods of assembly; short read-only, long read-only or a hybrid method utilizing both short and long reads. The user can generate an input CSV file containing multiple isolate samples from different species, all of which can be fed into the Reads2Resistome pipeline under a user-specified assembly method. Pipeline output is generated for quality control, assembly and annotation for each isolate. The pipeline is executable on both Mac and Linux operating systems and is well-suited for institutions and organizations which maintain, or have access to, a high-performance cluster for the analysis of “big data.”

Results from our case study indicated that obtaining a highly contiguous genome assembly with robust gene annotation, bacteriophage identification, and resistome characterization is best obtained under a hybrid assembly approach. While hybrid assembly is the most time-intensive assembly method, it produces the most complete annotated genome in our case study [23,24]. Long read-only assembly is able to produce a respectable genome length with high contiguity but falls short when annotating genomic features.

The ability to input multiple samples via the input CSV allows users to analyze hundreds of samples regardless of their identity. This prevents tedious bash scripting and collecting of various output files and ensures consistent and correct naming of all output files. Pipelines such as bacass from nf-core [25] provide genome assembly under short, long and hybrid approaches but do not offer resistome annotation. PRAP [26] and sraX [27] offer robust resistome analysis but do not provide genome assembly within the pipeline. Therefore, Reads2Resistome is the unique in providing a robust pipeline for genome assembly, genome annotation and resistome identification.

## Funding

This work has been supported by funding from the U.S. Department of Agriculture (USDA) Agricultural Research Service (ARS) Research Participation Program.

## Conflict of Interest

none declared.

## Notes

https://github.com/BioRRW/Reads2Resistome

### Competing Interest Statement

The authors have declared no competing interest.

## References

[1] T. Frieden, “Antibiotic resistance threats in the United States,” Centers Dis. Control Prev., p. 114, 2013, doi: CS239559-B.

[2] M. Su, S. W. Satola, and T. D. Read, “Genome-based prediction of bacterial antibiotic resistance,” Journal of Clinical Microbiology, vol. 57, no. 3. American Society for Microbiology, 01-Mar-2019, doi: 10.1128/JCM.01405-18.

[3] P. DI Tommaso, M. Chatzou, E. W. Floden, P. P. Barja, E. Palumbo, and C. Notredame, “Nextflow enables reproducible computational workflows,” Nature Biotechnology, vol. 35, no. 4. Nature Publishing Group, pp. 316–319, 11-Apr-2017, doi: 10.1038/nbt.3820.

[4] G. M. Kurtzer, V. Sochat, and M. W. Bauer, “Singularity: Scientific containers for mobility of compute,” PLoS One, vol. 12, no. 5, p. e0177459, May 2017, doi: 10.1371/journal.pone.0177459.

[5] A. M. Bolger, M. Lohse, and B. Usadel, “Trimmomatic: A flexible trimmer for Illumina sequence data,” Bioinformatics, vol. 30, no. 15, pp. 2114–2120, 2014, doi: 10.1093/bioinformatics/btu170.

[6] W. De Coster, S. D’hert, D. T. Schultz, M. Cruts, and C. Van Broeckhoven, “NanoPack: visualizing and processing long-read sequencing data,” doi: 10.1093/bioinformatics/bty149.

[7] R. R. Wick, L. M. Judd, C. L. Gorrie, and K. E. Holt, “Unicycler: Resolving bacterial genome assemblies from short and long sequencing reads,” PLOS Comput. Biol., vol. 13, no. 6, p. e1005595, Jun. 2017, doi: 10.1371/journal.pcbi.1005595.

[8] S. Nurk et al., “Assembling genomes and mini-metagenomes from highly chimeric reads,” Lect. Notes Comput. Sci. (including Subser. Lect. Notes Artif. Intell. Lect. Notes Bioinformatics), vol. 7821 LNBI, pp. 158–170, 2013, doi: 10.1007/978-3-642-37195-0_13.

[9] D. Antipov, A. Korobeynikov, J. S. McLean, and P. A. Pevzner, “HybridSPAdes: An algorithm for hybrid assembly of short and long reads,” Bioinformatics, vol. 32, no. 7, pp. 1009–1015, 2016, doi: 10.1093/bioinformatics/btv688.

[10] H. Li, “Minimap and miniasm: fast mapping and de novo assembly for noisy long sequences,” doi: 10.1093/bioinformatics/btw152.

[11] R. Vaser, I. Sović, N. Nagarajan, and M. Šikić, “Fast and accurate de novo genome assembly from long uncorrected reads,” Genome Res., vol. 27, no. 5, pp. 737–746, May 2017, doi: 10.1101/gr.214270.116.

[12] T. Seemann, “Prokka: Rapid prokaryotic genome annotation,” Bioinformatics, vol. 30, no. 14, pp. 2068–2069, Jul. 2014, doi: 10.1093/bioinformatics/btu153.

[13] T. Seemann, “ABRICATE,” Github, [Online]. Available: https://github.com/tseemann/abricate.

[14] M. Hunt, C. Newbold, M. Berriman, and T. D. Otto, “A comprehensive evaluation of assembly scaffolding tools,” Genome Biol., vol. 15, no. 3, p. R42, Mar. 2014, doi: 10.1186/gb-2014-15-3-r42.

[15] R. R. Wick, M. B. Schultz, J. Zobel, and K. E. Holt, “Bandage: interactive visualization of de novo genome assemblies,” doi: 10.1093/bioinformatics/btv383.

[16] S. K. Gupta et al., “ARG-annot, a new bioinformatic tool to discover antibiotic resistance genes in bacterial genomes,” Antimicrob. Agents Chemother., vol. 58, no. 1, pp. 212–220, Jan. 2014, doi: 10.1128/AAC.01310-13.

[17] B. P. Alcock et al., “CARD 2020: antibiotic resistome surveillance with the comprehensive antibiotic resistance database,” Nucleic Acids Res., vol. 48, no. D1, pp. D517–D525, Jan. 2020, doi: 10.1093/nar/gkz935.

[18] S. M. Lakin et al., “MEGARes: an antimicrobial resistance database for high throughput sequencing,” Database issue Publ. online, vol. 45, 2017, doi: 10.1093/nar/gkw1009.

[19] M. Feldgarden et al., “Validating the AMRFINder tool and resistance gene database by using antimicrobial resistance genotype-phenotype correlations in a collection of isolates,” Antimicrob. Agents Chemother., vol. 63, no. 11, 2019, doi: 10.1128/AAC.00483-19.

[20] A. Carattoli et al., “In Silico detection and typing of plasmids using plasmidfinder and plasmid multilocus sequence typing,” Antimicrob. Agents Chemother., vol. 58, no. 7, pp. 3895–3903, 2014, doi: 10.1128/AAC.02412-14.

[21] E. Zankari et al., “Identification of acquired antimicrobial resistance genes,” doi: 10.1093/jac/dks261.

[22] B. Liu, D. Zheng, Q. Jin, L. Chen, and J. Yang, “VFDB 2019: a comparative pathogenomic platform with an interactive web interface.,” Nucleic Acids Res., vol. 47, no. D1, pp. D687–D692, Jan. 2019, doi: 10.1093/nar/gky1080.

[23] N. De Maio et al., “Comparison of long-read sequencing technologies in the hybrid assembly of complex bacterial genomes,” Microb. Genomics, vol. 5, no. 9, 2019, doi: 10.1099/mgen.0.000294.

[24] S. Goldstein, L. Beka, J. Graf, and J. L. Klassen, “Evaluation of strategies for the assembly of diverse bacterial genomes using MinION long-read sequencing,” BMC Genomics, vol. 20, no. 1, p. 23, Jan. 2019, doi: 10.1186/s12864-018-5381-7.

[25] P. A. Ewels et al., “nf-core: Community curated bioinformatics pipelines,” bioRxiv, p. 610741, Apr. 2019, doi: 10.1101/610741.

[26] Y. He et al., “PRAP: Pan Resistome analysis pipeline,” BMC Bioinformatics, vol. 21, no. 1, p. 20, Jan. 2020, doi: 10.1186/s12859-019-3335-y.

[27] L. G. Panunzi, “sraX: A Novel Comprehensive Resistome Analysis Tool,” Front. Microbiol., vol. 11, p. 52, Feb. 2020, doi: 10.3389/fmicb.2020.00052.

